# Molecular Mechanisms Governing Shade Responses in Maize

**DOI:** 10.1101/397596

**Authors:** Qingbiao Shi, Fanying Kong, Haisen Zhang, Yu’e Jiang, Siqi Heng, Ran Liang, Jisheng Liu, Xiaoduo Lu, Pinghua Li, Gang Li

## Abstract

Light is one of the most important environmental factors affecting plant growth and development. Plants use shade avoidance and shade tolerance strategies to adjust their growth and development thus increase their success in the competition for incoming light. To investigate the mechanism of shade responses in maize (*Zea mays*), we examined the anatomical and transcriptional dynamics of the early shade response in seedlings of the B73 inbred line. Transcriptome analysis identified 912 differentially expressed genes, including genes involved in light signaling, auxin responses, and cell elongation pathways. Grouping transcription factor family genes and performing enrichment analysis identified multiple types of transcription factors that are differentially regulated by shade and predicted putative core genes responsible for regulating shade avoidance syndrome. For functional tests, we ectopically over-expressed *ZmHB53*, a type II HD-ZIP transcription factor gene significantly induced by shade, in *Arabidopsis thaliana*. Transgenic Arabidopsis plants overexpressing *ZmHB53* exhibited narrower leaves, earlier flowering, and enhanced expression of shade-responsive genes, suggesting that ZmHB53 participates in the regulation of shade responses in maize. This study increases our understanding of the regulatory network of the shade response in maize and provides a useful resource for maize genetics and breeding.

**Highlight:** Our findings not only increase the understanding of the regulatory network of the shade avoidance in maize, and also provide a useful resource for maize genetics and breeding.

## Introduction

Light plays a fundamental role in plant growth and development. Increasing the planting density of crops, particularly grasses such as maize (*Zea mays*), to increase yield is a common practice in modern agriculture. However, under high-density cultivation, light, water, and nutrients limit plant growth and seed production. Blue and red wavelengths light are preferentially absorbed by photosynthetic pigments of the upper leaves of the canopy for photosynthesis, resulting in a reduction of photosynthetically active radiation (PAR), and low ratio of red to far-red (R:FR) light in the lower leaves. In most plant species, the reduction of PAR, low R:FR and low blue light act as shade signals to induce shade avoidance syndrome (SAS), including elongation of stems and petioles and inhibition of the outgrowth of axillary buds, thus allowing plants to reach light and shade their neighbors (Keuskamp et al., 2010; Sharwood et al., 2014; Ballaré et al., 2017; Pignon et al., 2017). Long-term shade treatments lead to severe SAS and significantly decrease seed production (Casal, 2013); for example, maize grain yield may be reduced by up to 60% by long-term shade treatment (Cui et al., 2015). Therefore, understanding shade avoidance responses and improving plant success in the competition for light, without decreasing yields, are important goals to improve high-density planting of crops (Maddonni et al., 2001; Page et al., 2010).

The molecular network regulating SAS has been well documented in *Arabidopsis thaliana*. Various shade signals are primarily perceived by photoreceptors, including phytochromes and cryptochromes. Under high R:FR light, phytochromes (mostly phyB in Arabidopsis) enter the nucleus in the active form (far red-absorbing form, Pfr) and regulate numerous downstream genes, thereby suppressing the shade response (Kircher et al., 1999; Franklin, 2008; Chen et al., 2011). Low R:FR increases the ratio of inactive phyB (red-absorbing form, Pr) in the cytosol, thus releasing the inhibition of downstream signaling components and promoting the shade response (Kircher et al., 1999). The Arabidopsis *phyb* mutant and the maize *phyb1 phyb2* double mutant exhibit a constitutive SAS phenotype, including slender petioles, leaves, and accelerated stem elongation (Robson et al., 1993; Sheehan et al., 2007). Branch formation is also inhibited in the early development of *phyb* mutants in sorghum (Kebrom et al., 2016).

PHYTOCHROME INTERACTING FACTORS (PIFs) act as important downstream signal transduction components of phytochromes and play a key role in SAS (Castillon et al., 2007; Leivar et al., 2011). Arabidopsis plants overexpressing *PIF4, PIF5*, and *PIF7* exhibit constitutive SAS under high R:FR conditions (Lorrain et al., 2008; Li et al., 2012). Consistent with this, the *pifq* (*pif1 pif3 pif4 pif5*) quadruple mutant and *pif7* mutants show short petioles and a reduced response to shade (Leivar et al., 2008; Li et al., 2012). Overexpressing *ZmPIF4* in Arabidopsis also produces constitutive SAS (Shi et al., 2018). Analyses of genome-wide downstream targets revealed that PIFs directly target hundreds of growth-promoting genes, such as *Aux/IAA* (*IAA19, IAA29*), *YUCCA* (*YUC2, YUC5, YUC8, YUC9*), *EXPANSINS* (*EXPA1, EXPB1*) and *XYLOGLUCAN ENDOTRANSGLYCOSYLASE/HYDROLASE* (*XTH15, XTH33*) (Zhang et al., 2013; Pfeiffer et al., 2014). In addition, the contents and sensitivities of free auxin, gibberellin (GA) and brassinosteroids were rapidly induced by shade treatments, thus promoting cell elongation in Arabidopsis (Bou-Torrent et al., 2014), bean (Beall et al., 1996) and sunflower (Kurepin et al., 2007).

In Arabidopsis, shade treatments rapidly induce the expression of many transcription factor genes, including *LONG HYPOCOTYL IN FAR-RED* (*HFR1*), *PHYTOCHROME RAPIDLY REGULATED GENE 1* (*PAR1*), *PAR2*, and *PIF3-LIKE1* (*PIL1*), which encode basic helix-loop-helix (bHLH) type transcription factors that negatively regulate PIF activities through physical interactions, thereby preventing an exaggerated shade response (Roigvillanova et al., 2006; Hornitschek et al., 2009; Hornitschek et al., 2012). Additionally, multiple homeodomain leucine zipper (HD-ZIP) and B-box (BBX) type transcription factors function in the shade response (Sorin et al., 2009; Gangappa et al., 2014).

In maize, although some of the early shade-responsive genes have been identified (Wang et al., 2016), their physiological functions and underlying mechanisms remain largely unknown. Here, we combined cytological and transcriptomic analysis with functional testing to investigate the anatomical and transcriptional dynamics of SAS in maize seedlings and predict the core responsive genes involved in the regulation of SAS.

## Materials and Methods

### Plant materials and growth conditions

Seedlings of maize inbred lines were grown in growth chambers under a 12-hour light/12-hour dark cycle at 180 μmol/m^2^/s of light intensity at 25 °C. For short-term simulated shade treatment, seedlings of B73 (V3 stage) were transferred from white light to constant FR light (10.52 μmol/m^2^/s) for 0, 1, and 3 h, followed by constant R light (50 μmol/m^2^/s) for 1 h and then used for qPCR and RNA-seq assays. For long-term simulated shade treatment (Figure 1 and S1), various inbred lines were grown under white light (65.6 μmol/m^2^/s) supplied with FR light (10.52 μmol/m^2^/s, low R:FR), or R light (50.0 μmol/m^2^/s, high R:FR) after seed germination. After shade treatment, scanning electron microscopy (SEM) of sheath and leaf blade tissues were performed as previously described (Kong et al., 2017).

**Fig. 1.**
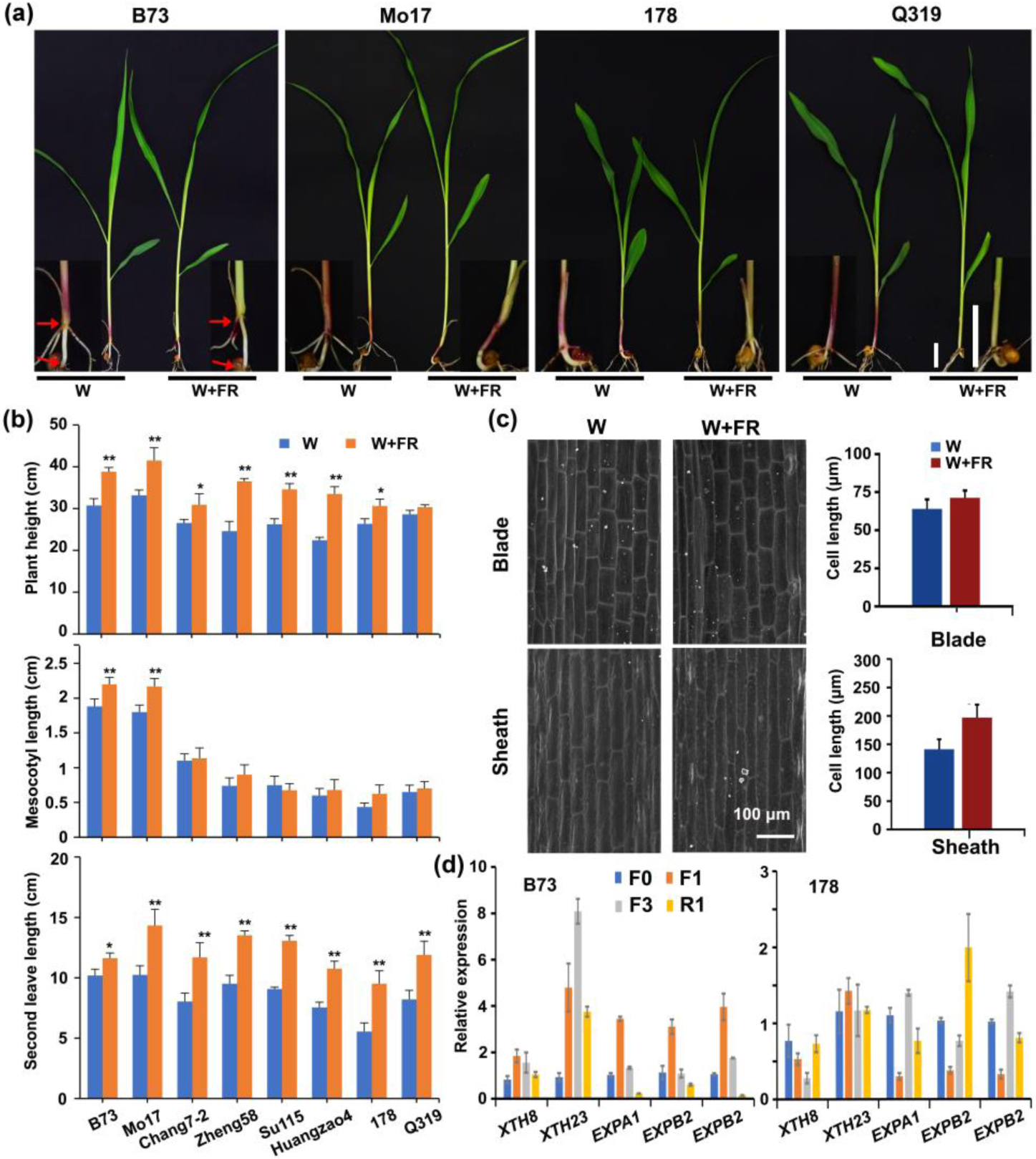
The phenotype of maize seedlings grown under high or low R:FR conditions. (a) The phenotypes of B73, Mo17, 178, and Q319 seedlings grown under high R:FR (13.3) and low R: FR light (0.19). Scale bar, 3 cm. (b) The plant height, mesocotyl length, and second-leaf length of different inbred lines. Data represent the mean and SD of at least 30 seedlings. *P < 0.05, **P < 0.01 (c) SEM and cell length analysis of the blade and sheath tissues of inbred B73 grown under high or low R:FR conditions. Data represent the mean and SD of at least 100 cells. *P < 0.05, **P < 0.01 (d) RT-qPCR analysis of the transcription level of cell elongation-related genes in B73 and 178 treated with far red light for 0 h (F0), 1 h (F1), and 3 h (F3), and then with red light for 1 h (R1), respectively. Data are means and SD of three independent biological replicates.

The *Arabidopsis thaliana* wild-type control plants used in this study were ecotype Columbia-0 (Col-0). The seeds were surface-sterilized with 20% bleach for 20 min and washed four times with sterile ddH_2_O. After being stratified for two days at 4 °C, the seeds were germinated on germination medium (GM) plates.

### RT-qPCR

Total RNA was extracted using an Ultrapure RNA kit (CWBIO, Beijing). The reverse-transcription reactions were performed using an AMV reverse transcriptase (Fermentas). The RT-qPCR was performed on a 7500 Fast Real-Time PCR machine (ABI) using SYBR Real Master Mix (Tiangen, Beijing). Primers used for RT-qPCR are listed in Table S3.

### RNA-seq analysis

The cDNA library construction, sequencing, and data analyses were performed as described previously (Kong et al., 2017). The maize B73 reference genome (AGPv3.22) were used for mapping the reads. The Cufflinks 2.2.1 package was used to calculate the gene expression levels with the parameter of reads per kilobase per million mapped reads (RPKM) and detect differentially expressed genes (DEGs) using default parameters. The false discovery rate (FDR) was used to determine the threshold of the p-value in multiple tests. A threshold of FDR ⩽0.05 and a fold change ⩾ 2 were used to judge the significance of differences in gene expression.

### Cluster and functional enrichment analysis

DEGs that were commonly expressed under both FR light and after re-exposure to R light (Dataset S2) and the expressed transcription factor genes were subjected to cluster analysis (Dataset S1). The RPKM values (normalized to the maximum of all RPKM values of the gene in B73 seedlings treated with FR light for 0 h, 1 h or 3 h, followed by R light for 1 h) were subjected to cluster analysis using the K-Means Support (KMS) module in the MultiExperiment Viewer (MEV) program.

### Plasmid construction and generation of transgenic Arabidopsis plants

To generate transgenic *ZmHB53-OE* lines in the Arabidopsis Col-0 background, the coding region of *ZmHB53* was PCR-amplified from cDNA of inbred line B73 using the primers pair *ZmHB53-F* and *ZmHB53-R* (Table S3). Then, *ZmHB53* fragment was inserted into the *BamH*I and *Xba*I digested *pPZP211-35Spro∷3FLAG* binary vector (Ma et al., 2017) to produce *35Spro∷ZmHB53-3FLAG*. More than 20 independent transgenic lines were selected and verified by RT-qPCR, followed by immunoblot analysis as described previously (Ma et al., 2016).

## Results

### Low R:FR induces the SAS in maize seedlings

To investigate the effects of shade on maize growth, seedlings of various inbred lines were grown under white light supplied with FR (R:FR ratio 0.19) or R (R:FR ratio13.3) conditions. After simulated shade treatment, the mesocotyl length, leaf length, and plant height significantly increased in inbred lines B73, Mo17, Huangzao4, Zheng58 and Su115, compared to plants under high R:FR conditions (Fig. 1a, 1b, S1). Mesocotyl length increased more strongly in inbred B73 (by 17%) and Mo17 (20%), compared with the other inbred lines (Fig.1b, S1). Moreover, the inbred lines 178 and Q319 were less responsive to simulated shade-induced elongation of mesocotyls and plant height, compared to the inbred lines B73 and Mo17 (Fig. 1). In addition, anthocyanin accumulation obviously decreased in the base region of the sheath in all the tested inbred lines under low R:FR conditions, compared with control plants grown under high R:FR conditions (Fig.1a, S1a).

To investigate the effects of simulated shade on cell elongation in B73, we observed the epidermal cells of the leaf blade and sheath by SEM. As shown in Figure 1c, cell elongation in the leaf blade increased slightly, while cell elongation in the sheath increased substantially after shade treatment. To further explore the effects of supplemental FR on cell elongation, we examined the transcript levels of cell elongation-related genes in V3 stage B73 and 178 seedlings that were transferred from white light to FR light for 1 and 3 h, followed by 1 h in red light. The transcript levels of *XTH8, XTH23*, and *EXPB2* were significantly induced by FR light and repressed by R light in inbred B73, but showed no obvious change in 178 (Fig. 1d), consistent with its reduced sensitivity to simulated shade treatment in Figure 1a-b.

### Generation and analysis of RNA-seq data for treated plants

To gain insight into the molecular regulatory mechanism of the shade response in maize, we conducted global RNA-seq of B73 seedlings at the V3 stage treated with FR light for 0 h (F0), 1 h (F1), or 3 h (F3), followed by R light for 1 h (R1) (short-term shading treatment). Using paired-end Illumina sequencing, we generated sequences from eight libraries (four time points with two biological replicates), producing approximately 1.9 billion high-quality reads, 95% of which uniquely mapped to the B73 reference genome, version 3. The distribution of reads was 75.8% in exons, 9.3% in introns, and 11.2% in intergenic genomic regions (Table S1). Comparisons of the biological replicates showed that their expression values were highly correlated (average R^2^ = 0.963, Fig. S2), indicating that the results of biological replicates in this study are highly reproducible. To reduce the influence of transcriptional noise, genes from the B73 filtered gene set (FGS) were included for analysis only if their RPKM values were ⩾1. In total, 22,479 genes were expressed under at least one condition, including 18,968 (84.5%) genes commonly expressed among all four conditions (Fig. 2a, Dataset S1).

**Fig. 2.**
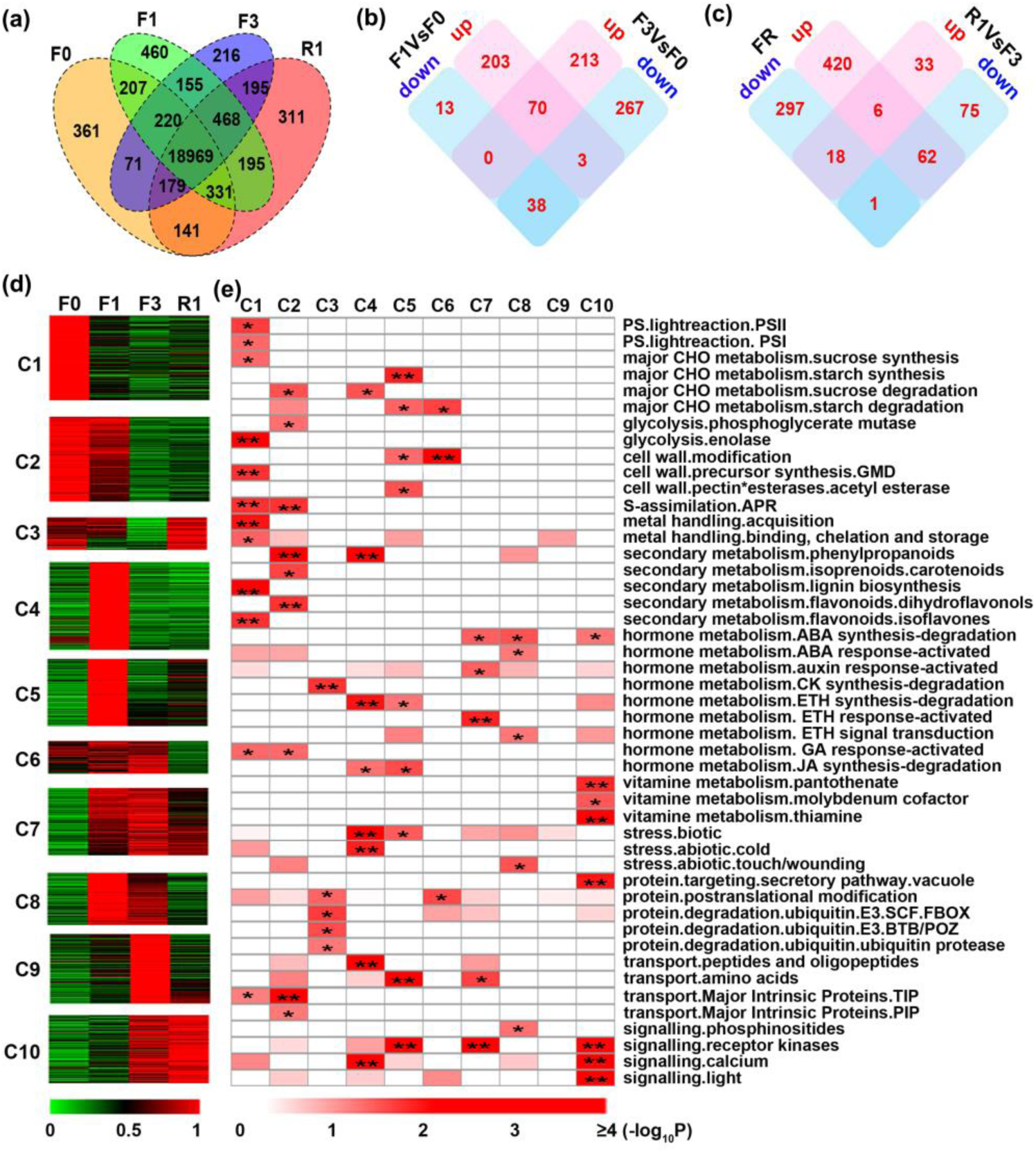
Transcriptome analysis of maize seedling responses to simulated shade. (a) Venn diagram of the numbers of the expressed genes in B73 seedlings treated with far red light for 0 h (F0), 1 h (F1), and 3 h (F3), and then with red light for 1 h (R1). (b) Venn diagram of the numbers of DEGs between F1 and F0 (F1 vs. F0) and F3 and F0 (F3 vs. F0), respectively. (c) Venn diagram of the numbers of the FR DEGs and DEGs between R1 and F3 (R1 vs. F3). FR DEGs refer to the DEGs of F1 vs. F0 and F3 vs. F0, excluding the 3 genes showing different trends. (d) Ten expression clusters of DEGs (C1–C10), ordered according to the time points of their peak expression. For each gene, the normalized values are shown. (e) Mapman functional enrichment analysis of DEGs. Fisher’s exact test was used to determine whether a functional category was enriched. *, *q* < 0.05; ** *q* < 0.01

To verify the quality of RNA-seq data, we performed RT-qPCR analyses of 48 transcripts, revealing a high correlation (R^2^ = 0.587) between the RNA-seq and RT-qPCR data (Fig. S3b). As expected, *ZmphyA1, ZmphyB1, ZmphyB2, ZmphyC1*, and *ZmphyC2* were significantly induced by FR (Fig. S3a). *ZmHY5* was strongly down-regulated by FR. Additionally, multiple genes encoding proteins involved in the light reactions in photosynthesis, such as *ZmLHCB1, ZmPSBA, ZmPSBQ*, and *ZmPSB28* were downregulated by FR (Fig. S3a).

Further, we identified 327, 591, and 195 DEGs between F0 and F1 (F1 vs. F0), F0 and F3 (F3 vs. F0) and F3 and R1 (R1 vs. F3), respectively (Fig. 2b and c, Dataset S2). Among these, 111 genes were common between F1 vs. F0, and F3 vs. F0, including three genes showing opposite expression patterns. Therefore, after excluding these three oppositely expressed genes, a total of 804 FR-regulated DEGs were identified (Fig. 2b). Interestingly, among the 87 common DEGs between FR-regulated, and red-regulated (R1 vs. F3), 80 (92%) genes showed opposite expression patterns, suggesting that most of the effects of FR light on gene expression can be reversed by subsequent treatment with R light (Fig. 2c). All these DEGs (912 genes) were selected as putative conserved genes important for the SAS in maize.

### Dynamics of gene expression during the SAS in maize

To better understand the regulatory network of the SAS in maize, we further grouped these 912 genes into 10 clusters (C1–C10) based on their expression patterns and then subjected to MapMan functional enrichment analysis (Fig. 2c). Among clusters (C1–C3) with reduced expression by FR, the most highly enriched categories included genes encoding proteins that mediate the light reactions, sucrose synthesis, and secondary metabolic pathways (Fig. 2e). For example, most of the anthocyanin biosynthesis related genes, including *CHS, CHI, F3H, DFR*, and *ANS* were highly downregulated by FR (Fig. S4), consistent with the reduced anthocyanin accumulation in shade-treated plants (Fig. 1a).

Among clusters of genes whose expression was induced by FR (C4–C10), the most highly enriched categories included genes encoding proteins involved in cell wall modification, degradation of starch and sucrose, hormone metabolism, and various signal pathways, suggesting they might play important roles in early responses to shade in maize. For example, genes related to auxin biosynthesis (*e.g., ZmYUC5* and *ZmTAA1*), and ethylene signal transduction (*e.g., ZmERF7*) were significantly induced by FR and downregulated by subsequent R treatment (Fig. 2e, S5). Interestingly, most of the alterations in expression (up- or downregulation) induced by FR were reversed by subsequent R treatment in most clusters, except for C10, which was enriched for genes involved in vitamin metabolism, protein targeting, and signaling. All these results are consistent with the regulatory network controlling the SAS in Arabidopsis (Li et al., 2011), suggesting that the regulatory mechanism of the SAS is evolutionarily conserved between monocots and dicots.

### Transcription factors play important roles in the maize SAS

Of the 3,316 maize transcription factor genes identified in the Plant Transcription Factor Database (http://planttfdb.cbi.pku.edu.cn/), 1,353 (41%) were commonly expressed under all four treatment conditions (Dataset S1). These genes were further classified into five groups based on their expression patterns (G1–G5, including 262, 212, 450, 191, and 238 genes, respectively; Fig. 3a). Shade-downregulated transcription factors were grouped into G1 and G2, including the HD-ZIP (21/43 expressed HD-ZIPs were included in G1 and G2) and MYB (44/124) transcription factors (Fig. 3b). Early shade-induced transcription factors were grouped into G3, which was significantly enriched for bHLH (41/109), ERF (46/92) and GRAS (21/48) family members (Fig. 3b, Dataset S1). Some bHLH family genes, including members of the PIF sub-family (*ZmPIF3, ZmPIF5*, and *PIF-like*) were rapidly induced by FR treatment. In addition, atypical *PIF* family genes, including *ZmHFR1, ZmPAR1, ZmPAR2* and *ZmPIL1*, were rapidly induced by shade and might play a negative role in the SAS (Dataset S1). Late shade-induced transcription factors were grouped in G4 and G5, and were significantly enriched for ARF (15/24) and HB/other (9/15) family members.

**Fig. 3.**
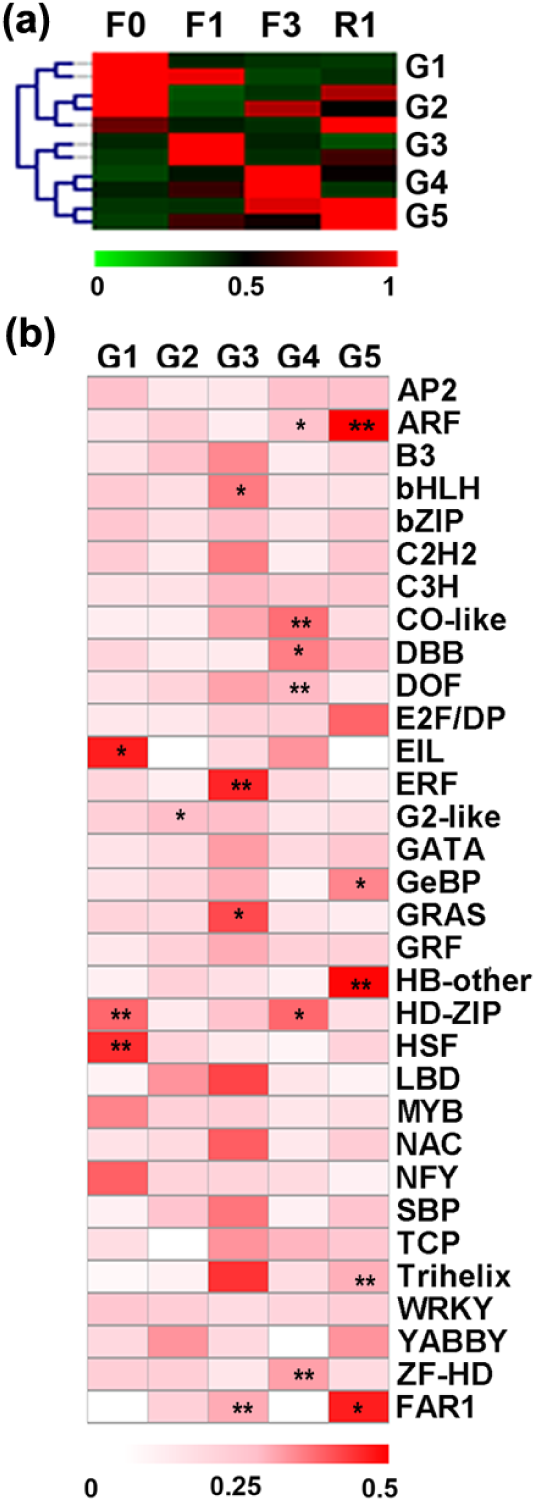
Transcription factor family enrichment analysis. (a) Five expression groups (G1–G5) of the expressed transcription factors. (b) Transcription factor family enrichment analysis. The values shown are the number of transcription factor family members classified in a cluster: the total number of transcription factor family members. Fisher’s exact test was used to determine whether a transcription factor family was enriched. *, *q* < 0.05; ** *q* < 0.01.

### Core genes involved in regulating the SAS

To identify the important regulators of the shade response, we first listed the 226 genes overlapping in our DEG list (912, Dataset S2) and Wang’s DEG list (1105, Wang et al., 2016), eliminated the photosynthesis, secondary metabolism, stress, nucleotide metabolism, and function unknown genes from this list, added three genes, *ZmGT1* (*Grassy tillers1*), *ZmTB1* (*Teosinte branched1*), and *ZmVT2*, which have already been shown to play important roles in maize SAS (Doebley et al., 1997; Sheehan et al., 2007; Phillips et al., 20011; Whipple et al., 2011), and identified 93 core genes for the shade response (Table1, S2). Most of these genes were significantly regulated by shade treatment. In addition to *ZmGT1, ZmTB1*, and *ZmVT2, ZmphyB1* and *ZmphyB2* have also been proved to participate in maize SAS (Sheehan et al., 2007). The other core shade-responsive genes have not previously been shown to directly regulate the SAS in maize, but are related to light signaling, hormone metabolism and signal transduction, regulation of transcription, cell wall modification, protein metabolism and so on (Table 1). For example, multiple plant hormone-related genes including *IAAs, SAURs* and *GH3.1, GA1, GA5, GA2ox1, GA2ox8, CKX6, ACO1, EIN4* and *ERFs* were identified, suggesting that they may play crucial roles in the SAS in maize (Table 1). Interestingly, we identified 7 BBXs as core genes for SAS regulation (Table 1, S2). For example, *ZmBBX20* was upregulated 2.9-fold (F1 vs. F0) and 12.7-fold (F3 vs. F0) in response to shade treatment in the current study, and 2.5-fold (1 h vs. 0h), 2.1-fold (3 h vs. 0 h) and 2.9-fold (6 h vs. 0 h) in the previous study (Wang et al., 2016).

**Table 1.**
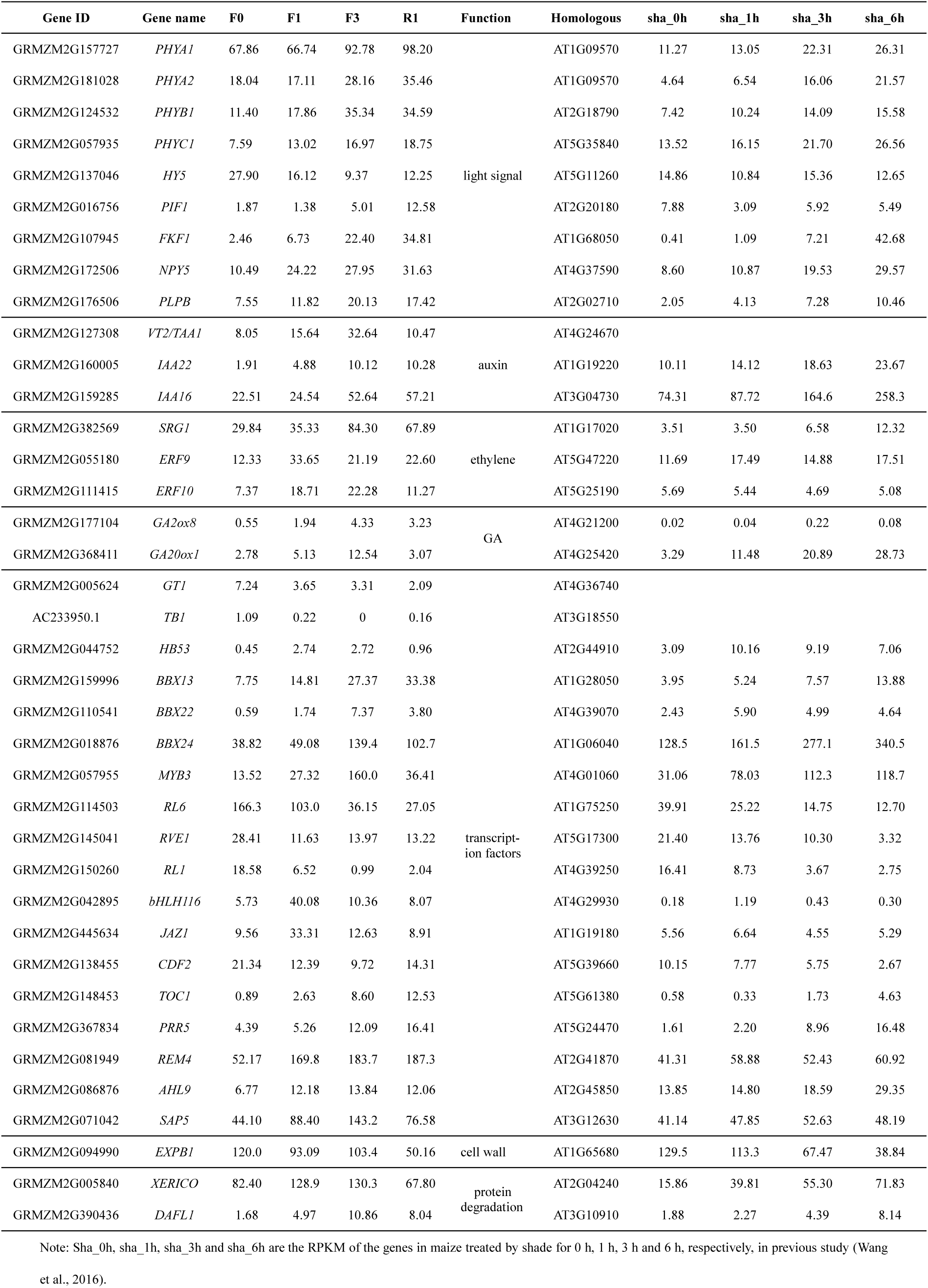
Core responsible genes involved in regulating the SAS in maize.

### ZmHD-ZIP proteins act as regulators of the SAS

Research in Arabidopsis has shown that HD-ZIP transcription factors modulate the SAS (Sorin et al. 2009). In our RNA-seq data, many HD-ZIP family genes were up- or downregulated by shade, therefore, this transcription factor family was selected for further analysis (Fig. 4a). Phylogenetic analysis of this family genes in Arabidopsis and maize revealed that these genes were classified into the I, II, III and IV subfamilies (Fig. 4b). Interestingly, one-third of type I HD-ZIP and all the type II HD-ZIP genes were up regulated, while two-thirds type I HD-ZIP genes were down regulated by FR (Fig. 4a), indicating that various members of this transcription factor family (for example type I and II HD-ZIP genes) might play opposite roles in the shade response. Consistent with the results of RNA-seq, qPCR analyses revealed that type II HD-ZIPs, including *ZmHB4, ZmHB53, ZmHB59, ZmHB78*, and *ZmHB86*, were strongly induced by FR, and subsequently suppressed by R light; by contrast, type I HD-ZIPs, such as *ZmHB34, ZmHB66*, and *ZmHB70*, were slightly reduced by FR and induced by R light (Fig. 4c). These results demonstrate that both the type I and II HD-ZIP subfamily members might play more important roles in the responses to shade signals.

**Fig. 4.**
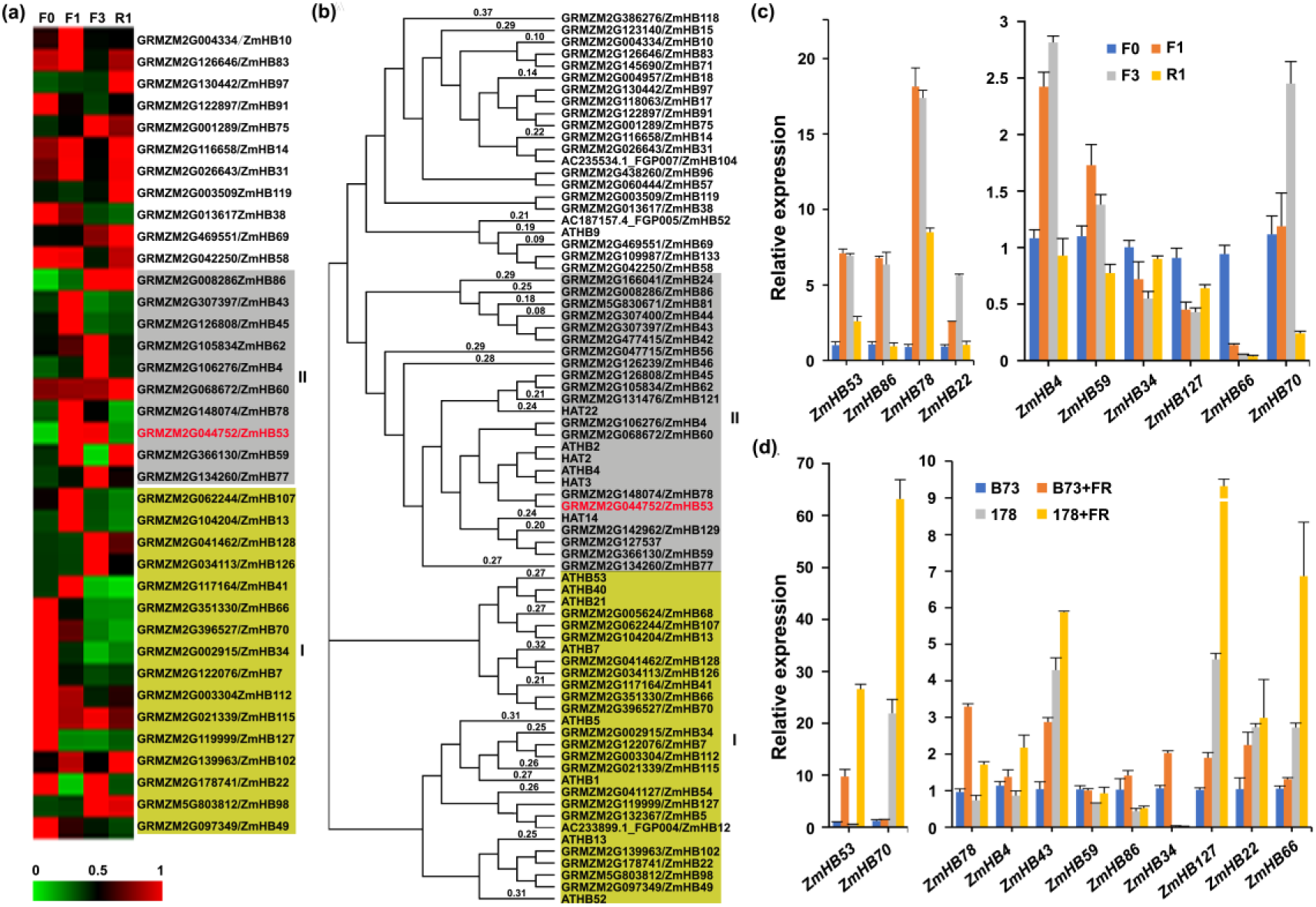
Expression pattern analysis of HD-ZIP members in shade response. (a) Heat map representation of the expression patterns of *HD-ZIPs*. For each gene, the value shown is the RPKM value normalized by the maximum values of all RPKM values of the gene in F0, F1, F3 and R1. The gene in gray background belong to type II *HD-ZIPs*, while in yellow background belong to type I *HD-ZIPs.* (b) Phylogenetic tree of selected HD-ZIP family proteins in *Zea mays* (Zm) and *Arabidopsis thaliana* (At). The neighbor-joining method was used to construct the phylogenetic tree. (c) RT-qPCR analyses revealed that selected HD-ZIP family genes were rapidly induced or reduced by far-red or red light. Three-leaf stage seedling plants of maize inbred line B73 were used to harvest second leaves, and then used to perform RT-qPCR analysis. *Actin* was used as an internal control for RT-qPCR analysis. Data are means and SD of three independent biological replicates. (d) RT-qPCR analyses of the expression of selected HD-ZIP family genes in B73 and 178 lines treated by white or FR for 1h.

We further measured the expression levels of these HD-ZIP genes in the B73 and 178 inbred lines under simulated shade conditions. As shown in Figure 4d, *ZmHB43, ZmHB53, ZmHB78* and *ZmHB127* were induced by shade in both inbred B73 and 178, while *ZmHB34, ZmHB66*, and *ZmHB70* showed opposite expression patterns in these inbred lines under shade treatment, suggesting these genes may contribute to the differential response to shade between the B73 and 178 inbred lines.

### ZmHB53 can affect the shade response in maize

To further investigate the roles of HD-ZIPs in the shade response, we focused on *ZmHB53* (GRMZM2G044752), a homolog of *ATHB4* which is essential in shade response and leaf development in Arabidopsis (Sorin et al. 2009). To investigate whether *ZmHB53* affects leaf morphogenesis and shade responses, we overexpressed a FLAG-tagged version of *ZmHB53* (*ZmHB53-3Flag*) under the control of the constitutive 35S promoter in the Arabidopsis Col-0 background. Three independents transgenic *ZmHB53* overexpression lines (*OE5, OE6*, and *OE8*) were selected based RT-qPCR and western-blotting, and then subjected to further physiological analysis (Fig. S5a).

All three lines exhibited a slight SAS, including narrow rosette leaves and early flowering time, compared with wild-type Col-0 plants under long-day (LD, 16-hour light/8-hour dark) conditions (Fig. 5a–b, 5d–e). Interestingly, the transgenic lines had more branches and reduced plant height compared to wild type at the mature stage, representing a lessened response to shade treatment, compared with wild-type control plants (Fig. 5c, 5e), indicating that *ZmHB53* can affect SAS in Arabidopsis via a complex regulatory mechanism. However, in seedlings grown under dark, white, and low R/FR light conditions, Arabidopsis *ZmHB53* overexpression lines showed no significant differences from wild-type control plants (Fig. S5), suggesting that *ZmHB53* mainly functions at the mature stage.

**Fig. 5.**
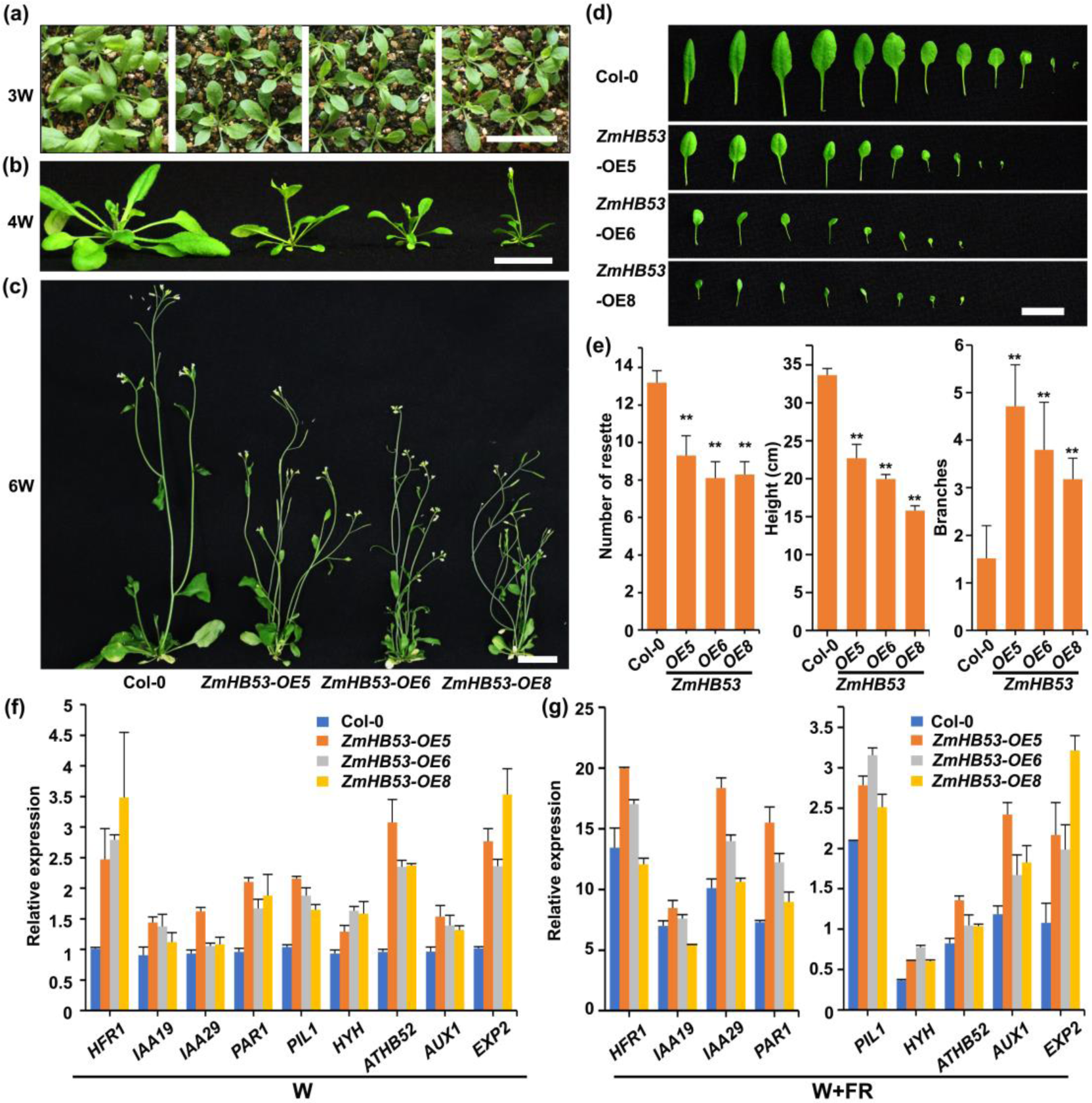
Overexpression of ZmHB53 in wild-type Arabidopsis Col-0 plants. (a-c), Phenotype of 3-, 4-, and 6-week-old Arabidopsis overexpressing *ZmHB53*, respectively. Scale bar, 2 cm. (d) Leaves of 4-week-old Arabidopsis overexpressing *ZmHB53.* Scale bar, 2 cm. (e) Quantification of rosette leaf number shown in B and D, plant height and number of branches shown in C. Scale bar, 2 cm. \**P <* 0.05; \**P <* 0.01; *n* = 20. (f-g) RT-qPCR analysis of the expression of selected shade-response genes in Arabidopsis overexpressing *ZmHB53*. Seven-day-old seedlings grown under LD (W, high R:FR; W+FR, low R:FR) conditions were used. *UBQ1* was used as the internal control. Data are means and SD of three replicates.

Next, we examined the transcript levels of genes that respond rapidly to shade treatment in the *ZmHB53-*overexpressing lines via RT-qPCR. Under white-light (R:FR 7.8) conditions, the transcript levels of well-known shade-responsive genes including *HFR1, PAR1, PIL1*, and *EXP2*, all significantly increased in the Arabidopsis *ZmHB53-* overexpressing lines, compared with wild-type Col-0 control plants (Fig. 5f). After simulated shade treatment (W+FR), the expression of *HFR1, PAR1*, and *PIL1*, were significantly upregulated in Col-0 and *ZmHB53* overexpression lines, compared with control plants under white light conditions (Fig. 5f–g). All these results indicate that overexpressing *ZmHB53* enhances the transcript levels of shade-response genes in Arabidopsis.

## Discussion

Light is one of the essential factors determining yield potential in the modern high-density cultivation of crop plants. In most plant species showing shade avoidance response, changes in light quantity and quality cause morphological responses including elongated stems and petioles, and more erect leaf angle; these responses increase leaf vertical inclination and help the plant compete for light (Zhu et al., 2014; Bongers et al., 2018). Here, we found that the maize inbred lines 178 and Q319 exhibited less-pronounced responses to simulated shade treatment, compared with inbred lines B73 and Mo17 (Fig.1a-b). B73 maize seedlings under simulated shade conditions showed typical symptoms of the SAS, such as elongated mesocotyls, stems, and leaves, and reduced accumulation of anthocyanin (Fig. 1a–b). Consistent with this, cytological, qPCR and RNA-seq analyses showed that simulated shade treatment induced the transcription levels of cell elongation-related genes and promoted cell elongation in leaf blades and sheaths (Fig. 1c–d).

Phytochrome signaling pathways play a conserved role in the low R:FR induced shade response in both maize and Arabidopsis (Lee et al., 2017). Arabidopsis PIF4, PIF5 and PIF7 act as the downstream signal transduction components of multiple photoreceptors (including phytochromes and cryptochromes) and play crucial roles in shade responses (Lorrain et al., 2008; Leivar et al., 2011; Li et al., 2012). Here, *ZmphyA1, ZmphyB1, ZmphyB2*, and five *PIF* family genes were all upregulated by FR, suggesting that they may play important roles in shade responses. Consistent with this, our previously study showed that the over-expression of *ZmPIF4* and *ZmPIF5* causes a constitutive shade avoidance response in Arabidopsis, indicating that they might play essential roles in shade responses in maize (Shi et al. 2018).

A reduction in the outgrowth of axillary buds is one of the typical morphological changes of the shade avoidance response. The Arabidopsis TCP (TEOSINTE BRANCHED 1, CYCLOIDEA, PCF) type transcription factor BRANCHED 1 (BRC1) directly binds to and activates the transcription of a group of HD-ZIP I transcription factor genes, including *HB21, HB40*, and *HB53*, thus preventing constitutive outgrowth of branches (Gonzalez-Grandio et al., 2017). Maize *TB1* is a homolog of *BRC1*, and negatively regulates the outgrowth of axillary buds (Doebley et al., 1997). Maize *GT1*, encoding an HD-ZIP I family member, is one of the downstream targets of TB1 and represses the outgrowth of lateral buds (Whipple et al., 2011). Therefore, it appears that the genetic module involving the BRC1/TB1 and HD-ZIP transcription factors is evolutionarily conserved in dicots and monocots, where it prevents branching under light-limiting conditions. Interestingly, the Arabidopsis *ZmHB53* (HD-ZIP II) overexpression lines showed more branches than the wild-type control plant, which contrasts with the phycological function of maize *GT1* (Fig. 5). Therefore, we hypothesized that HD-ZIP transcription factors, for example HD-ZIP I and II sub-family, may play negative and positive roles in regulating the outgrowth of axillary buds, respectively, like the functions of bHLH type transcription factors in SAS, such as the positive roles of PIF4 and PIF5, and the negative roles of HFR1, PAR1, and PAR2 in the shade response in Arabidopsis. This is also consistent with the opposite expression trends of type I and II HD-ZIP genes in response to shade in maize (Fig. 4a).

In summary, the monocotyledonous plant maize and the dicotyledonous plant Arabidopsis share a number of morphological and physiological responses in their shade responses. When plants are exposed to shade conditions, photoreceptor systems perceive a reduction of PAR, low ratio of R:FR, as well as low blue light, and subsequently activate a downstream network of various interacting transcriptional regulators and hormones to adjust plant growth and development to increase the plant’s ability to compete for light (Fig. S6). Based on this model, three different strategies could be developed to increase the ability of maize to compete for light and minimize the negative effects of the SAS. In the upper regulatory layer, one strategy could involve modulating the expression levels or activities of photoreceptor genes such as *ZmphyA1, ZmphyA2, ZmphyB1*, and *ZmphyB2*, as they directly respond to dynamic environmental light changes. At the middle regulatory layer, another strategy could modify the expression of important regulators of the SAS, such as *ZmPIF4, ZmPIF5, ZmHFR1* and *ZmHB53*. In the downstream regulatory layer, a third strategy could modify the expression levels of many SAS-related genes, including those directly involved in cell elongation, hormone synthesis, or signaling transduction, such as *ZmTAA1* and *ZmYUC5*. Finally, our study identified a core set of shade-responsive genes, which expands the regulatory network of shade responses and provides a useful resource for maize genetics and breeding.

## Supplemental Information

Supplemental information is available online.

Fig. S1. The phenotype of maize plants grown under high or low R:FR conditions.

Fig. S2. Correlation between biological replicates.

Fig. S3. Verification of RNA-seq results by RT-qPCR.

Fig. S4. Expression analysis of genes involved in anthocyanin biosynthesis in maize by RNA-seq.

Fig. S5. Identification of *ZmHB53* overexpression transgenic plants and the response of seedlings to shade.

Fig. S6. Model of the putative regulatory network of the early shade response in maize.

Table S1. RNA-seq data analysis.

Table S2. Core responsible genes involved in regulating the shade response in maize.

Table S3. Primers used in this study.

Dataset S1. Genes expressed during the shade response in maize.

Dataset S2. Differentially expressed genes during the shade response in maize.

## Acknowledgments

This work was supported by grants from the State Key Basic Research and Development Program of China (2014CB147301 and 2016YFD0101003), the National Key Research and Development Plan of China (2017YFD0301001 and 2018YFD0300603), the National Natural Science Foundation of China (31701434), Funds of Shandong “Double Tops” Program (to GL) and the State Key Laboratory of Crop Biology (DXKT201706).

